# Phosphoglycerate kinase is a central leverage point in Parkinson’s Disease driven neuronal metabolic deficits

**DOI:** 10.1101/2023.10.10.561760

**Authors:** Alexandros C. Kokotos, Aldana M. Antoniazzi, Santiago R. Unda, Myung Soo Ko, Daehun Park, David Eliezer, Michael G. Kaplitt, Pietro De Camilli, Timothy A. Ryan

## Abstract

Phosphoglycerate kinase 1 (PGK1), the first ATP producing glycolytic enzyme, has emerged as a therapeutic target for Parkinson’s Disease (PD), since a potential enhancer of its activity was reported to significantly lower PD risk. We carried out a suppressor screen of hypometabolic synaptic deficits and demonstrated that PGK1 is a rate limiting enzyme in nerve terminal ATP production. Increasing PGK1 expression in mid-brain dopamine neurons protected against hydroxy-dopamine driven striatal dopamine nerve terminal dysfunction *in-vivo* and modest changes in PGK1 activity dramatically suppressed hypometabolic synapse dysfunction *in vitro*. Furthermore, PGK1 is cross-regulated by PARK7 (DJ-1), a PD associated molecular chaperone, and synaptic deficits driven by PARK20 (Synaptojanin-1) can be reversed by increasing local synaptic PGK1 activity. These data indicate that nerve terminal bioenergetic deficits may underly a spectrum of PD susceptibilities and the identification of PGK1 as the limiting enzyme in axonal glycolysis provides a mechanistic underpinning for therapeutic protection.

- Suppressor screen identifies PGK1 as the rate limiting enzyme in axon metabolism
- PGK1 expression *in-vivo* protects against striatal dopaminergic axon degeneration
- Loss of DJ-1 impairs axonal ATP production
- PGK1 function is regulated by PARK7/DJ-1, and can reverse PARK20 synaptic deficits

## Introduction

The brain is a vulnerable metabolic organ that suffers acute functional decline when fuel delivery is compromised. We previously showed that CNS nerve terminals are one of the likely loci of metabolic vulnerability as they rely on efficient activity-dependent upregulation of ATP synthesis to sustain function, but failure to do so leads to abrupt synaptic collapse^1–3^. Reduced fuel delivery to the brain is correlated with the aging process and is considered an early predictor of eventual neurological dysfunction^4^, suggesting that as fuel delivery becomes compromised, synaptic function becomes increasingly vulnerable to genetic metabolic lesions. PD has long-been thought to be in part driven by the metabolic vulnerability of dopamine (DA) neurons of the substantia nigra compacta (SNc) as two of the earliest identified genetic drivers of PD, PARK2 (PARKIN) and PARK6 (PINK1), when mutated, compromise the integrity of mitochondria^5^, a central hub of cellular bioenergetic support. Furthermore, SNc DA neurons, which fire constantly due to their pacemaker activity, comprise extreme, elaborated axonal arbors, that in turn are thought to support up to 1 million DA release sites each in the human striatum, thus presenting a large metabolic burden^6^. In addition, the striatum, where SNc axons project, is characterized by low abundance of glucose^7^ and trophic factors. Many experimental animal models of PD specifically target bioenergetic pathways in DA neurons, as lesioning mitochondrial function in these neurons leads to their degeneration^8,9^.

The primary substrate for mitochondrial oxidative phosphorylation is pyruvate, the product of a 10-step enzymatic cascade in the cytosol that converts glucose into 2 molecules each of ATP and pyruvate. Several recent discoveries point to a critical but unexpected outsized role of the glycolytic enzyme PGK1 in protecting neurons against neurological impairment. PGK1, the first ATP producing enzyme in glycolysis, catalyzes the sixth step in this 10-step enzymatic cascade. A chemical screen of a subset of FDA-approved drugs capable of suppressing cell death identified Terazosin (TZ) as a weak activator of PGK1^10^. TZ was subsequently shown to confer significant protection in numerous models of PD (mouse, rat, drosophila and human iPSCs)^11^ implying that contrary to expectations, PGK1 activity is a critical modulator of glycolytic throughput. The clinical use of TZ for treatment of benign prostate hyperplasia (BPH) provided data for a retrospective analysis to determine the extent to which TZ might alter the disease emergence in humans. Remarkably, Simmering and colleagues concluded that, in humans, prolonged use of TZ, reduced the risk of developing PD by ∼ 37% compared to alternate BPH treatments^12^. Furthermore, PGK1 is part of the PARK12 susceptibility locus^13^, and certain PGK1 mutations in humans are characterized by early onset PD^14,15^.

These data all predict that PGK1 activity is a crucial leverage point in neuronal bioenergetic control. To test this idea, we developed a cellular approach in primary neurons to determine which, if any of the glycolytic enzymes in neurons, might be capable of improving metabolic resilience when faced with diminished fuel supply. Our data demonstrate that modest changes in PGK1 activity have dramatic impact on synapse function when operating under hypometabolic conditions and identify PARK7/DJ-1, the PD-associated molecular chaperone, as a critical PGK1 regulator. Increasing PGK1 abundance in vivo offered strong protection against striatal DA axon dysfunction and was able to reverse cellular phenotypes in primary cultured neurons harboring the PARK20 mutation. These data strongly support the idea that PGK1 activity serves as a critical lever arm in controlling nerve terminal function under hypometabolic conditions and suggests that hypometabolic deficits may underly a wide-spectrum of PD etiology^11,12,14–19^.

## Results

### Synaptic endurance: a measure of neuronal metabolic resilience

We previously established that presynaptic function in central nerve terminals is extremely sensitive to metabolic lesions, as they need local on-demand upregulation of ATP synthesis to meet their energetic demands during activity^1,3,20^. Inability to do so rapidly leads to failure in SV recycling^3,17^. To identify molecular pathways that might confer metabolic resilience we devised a synaptic endurance test based on previously established SV recycling assays carried out in primary dissociated hippocampal neurons. Here, neurons are subjected to repeated bouts of stimulation at regular intervals (Fig. 1A). With sufficient fuel (e.g. 5 mM glucose), SV recycling can be sustained through at least 10 rounds of 100 action potential (AP) bursts delivered at minute intervals as measured with vGlut-pHluorin (vGlut-pH) fluorescence (Fig. 1B, C). However, this synaptic endurance fails when extracellular glucose is lowered to 0.1 mM (Fig. 1B), as successive rounds of stimulation lead to a gradual slowing of the endocytic retrieval/reacidification process and a gradual increase in the post-stimulus fluorescence (Fig. 1C).

**Figure 1.**
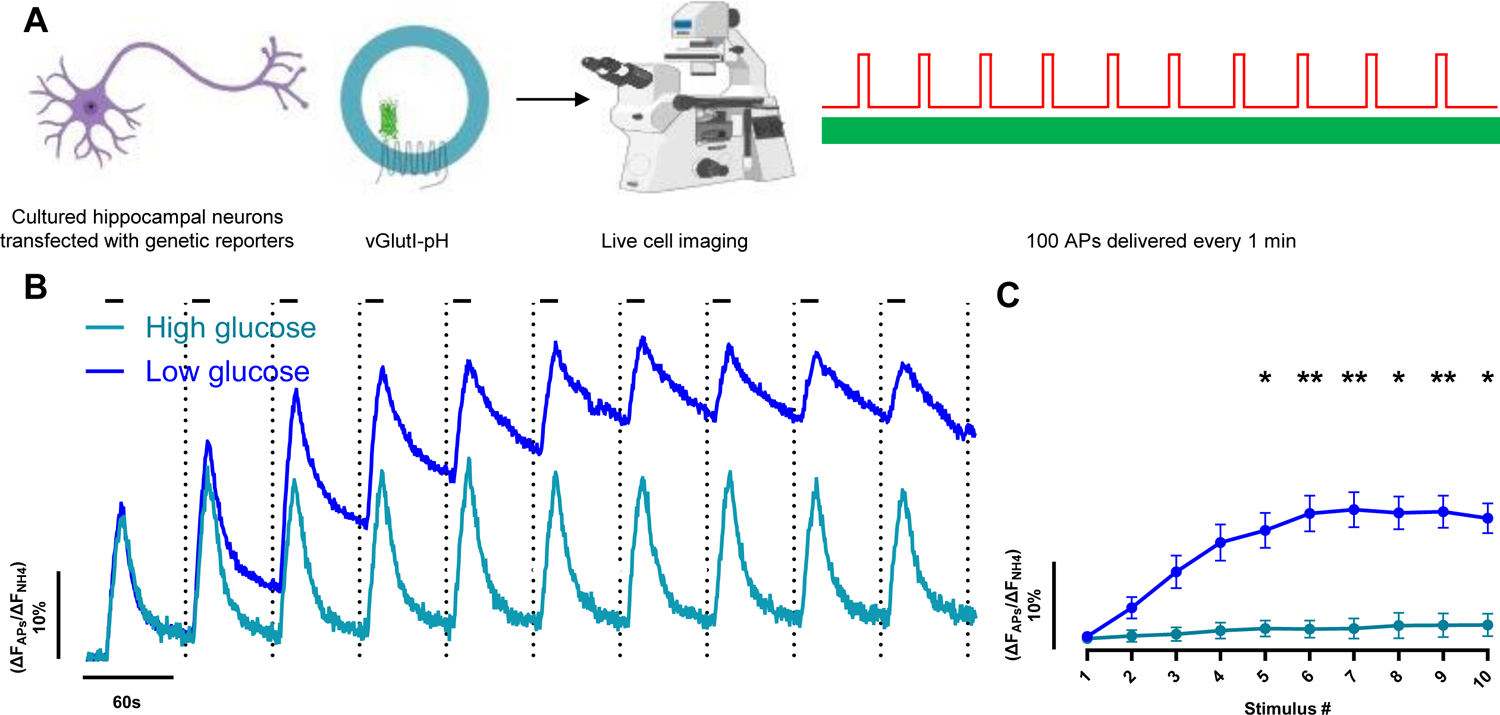
Hypometabolic synaptic endurance assay. **(A)** Assay schematic: Cultured primary hippocampal neurons expressing vGlutI-pH incubated in imaging media of interest for 5 min prior to ten bouts of AP firing (100 APs at 10 Hz, highlighted as black bars in the traces) delivered every minute. (**B**) Ensemble average vGlutI-pH responses are normalized to the maximal sensor fluorescence revealed by perfusion with 50 mM NH_4_Cl. With sufficient fuel (5 mM glucose, teal) efficient SV recycling persists for all rounds of stimulation but in low glucose (0.1 mM glucose, blue) SV recycling gradually slows, resulting in a net accumulation of fluorescence. (**C**) The ensemble average fraction of the fluorescence remaining 55 s post stimulation for each bout for high and low glucose conditions mean ± SEM for the data shown in (**B**), 5 mM glucose N=6, 0.1 mM glucose N=23 *p < 0.05, **p < 0.01 2-way ANOVA.

### A suppressor screen of hypometabolic synaptic failure identifies PGK1 as a rate limiting enzyme in presynaptic bioenergetics

The gradual synaptic failure under hypometabolic conditions allowed us to conduct genetic expression suppressor screen of glycolytic enzymes to see if any single one, when over-expressed, would allow synapses to sustain function under limited fuel availability. Of the enzymes evaluated (see methods), only one, PGK1, conferred significant hypometabolic resilience. PGK1, which, when overexpressed, was able to fully support synaptic function in 0.1 mM glucose, exhibiting robust SV recycling for all ten rounds of activity tested (Fig. 2A, B). This functional rescue was conducted using a HALO-tagged PGK1 allowing us to visualize the expressed protein distribution in live cells, confirming that the functional rescue was neuron specific. Quantitative immunofluorescence for PGK1 and synapsin showed that native PGK1, although expressed throughout the cell, was highly concentrated in nerve terminals (Fig. 2C, S1E, F). Quantitative comparison of presynaptic PGK1 immunofluorescence showed that, on average, our expression construct led to a ∼9-fold over-expression of this enzyme at nerve terminals (compared to non-transfected cells) that varied significantly across experiments (Fig. S1A, B, C,). Live cell labeling of the HALO moiety (Fig. 2D) allowed us to determine, on a cell-by-cell basis, the presynaptic abundance of PGK1-HALO and determine quantitatively how exogenous PGK1-HALO expression correlated with hypometabolic resilience (Fig. 2E). These data revealed that every cell overexpressing PGK1-HALO was able to confer metabolic resilience, even at the lowest expression level achieved, ∼ 2-fold (Fig. S1D). As expected, SV recycling was impaired in neurons expressing an shRNA targeting PGK1 (Fig. S2A, B, C).

**Figure 2.**
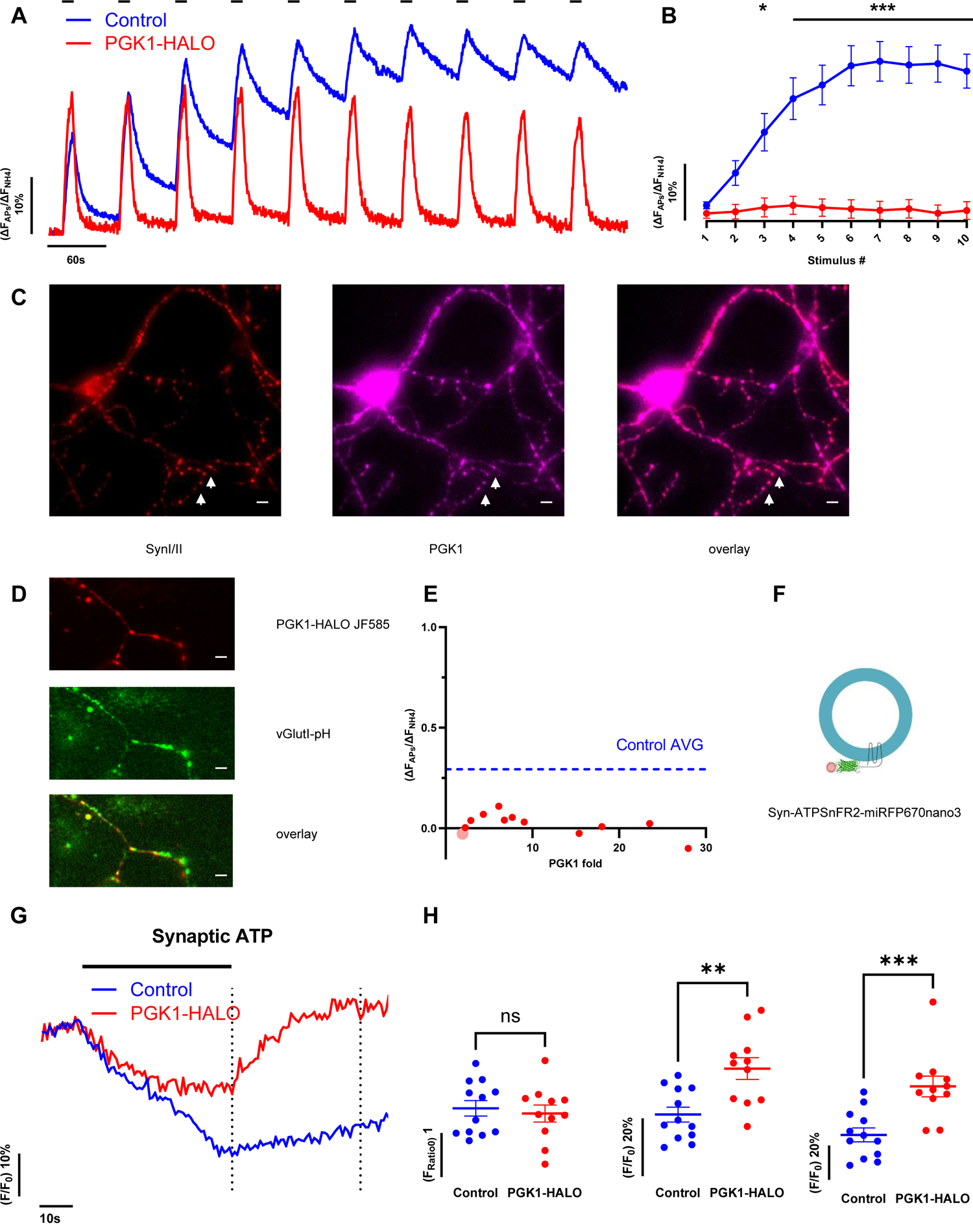
Synaptic PGK1 expression confers synaptic resilience under hypometabolic stress. **(A)** Ensemble average vGlutI-pH fluorescence in neurons expressing PGK1-HALO (red) shows that synaptic endurance is fully restored in 0.1 mM glucose compared to controls (blue). (**B**) The remaining fluorescence 55 s post stimulation after each train is plotted as mean ± SEM, control N=23, PGK1-HALO N=12 *p < 0.05, ***p < 0.001 2-way ANOVA. (**C**) PGK1 (magenta) and synapsin (red) immunofluorescence shows that PGK1 is present in nerve terminals (white arrows). (**D**) PGK1-HALO (labelled with JF585) also accumulates in nerve terminals (vGlutI-pH, visualized during NH_4_Cl application) scale bar in **C** and **D** 6 μm. (**E**) Synaptic endurance score, measured as the fluorescence signal of recovery after the 10^th^ round for each cell tested compared to the average synaptic PGK1-HALO expression normalized to non-transfected cells N=12. Dashed blue line shows the average synaptic endurance score for low glucose in control neurons after the 10^th^ round. (**F**) Schematic of the synapto-iATPSnFR2-miRFP670nano3 sensor used for synaptic ATP measurements. (**G**) Ensemble average synapto-iATPSnFR2-miRFP670nano3 traces for control (blue) and PGK1-HALO (red) transfected cells stimulated with 600 APs at 10 Hz in 0.1 mM glucose. PGK1-HALO neurons show a significant activity dependent upregulation of ATP synthesis following activity. (**H**) Comparison of absolute nerve terminal ATP values pre-stimulus (left) at the end of the stimulus (left dotted line in (PGK1-HALO)) (middle) and 35 s post-stimulus (right dotted line in **C**) (right) in control and PGK1-HALO expressing neurons, mean ± SEM indicated, control N=12, PGK1-Halo N=11. *p < 0.05, **p < 0.01 unpaired t-test.

The dramatic impact of PGK1 expression on hypometabolic synapse function strongly implies that increasing this enzyme’s abundance leads to a robust change in the kinetics of presynaptic ATP synthesis. To test this idea we carried out parallel measurements of presynaptic ATP dynamics using a next-generation genetically encoded ratiometric ATP reporter (synapto-iATPSnFR2.0-miRFP670nano3) targeted to nerve terminals^21^ (Fig. 2F). Under hypometabolic conditions, a sustained 60 s burst of 600 AP led to a 30% depletion of presynaptic ATP that only recovered minimally for the next 30 s (Fig. 2G, H). In neurons expressing PGK1-HALO, the initial prestimulus ATP values were similar to controls, however the decrease in ATP during the stimulus was blunted (decreasing by 14%), while in contrast to controls, ATP levels recovered rapidly over the next 30s. These data demonstrate that increased PGK1 abundance is sufficient to increase the kinetics of ATP production during demand, indicating that, following activity, PGK1 is a rate-limiting enzyme in nerve terminal glycolysis and that a modest change in abundance has dramatic functional consequences.

### PGK1 expression protects against dopaminergic axon degeneration in the striatum

Previous *in-vitro* analysis demonstrated that TZ could only very modestly accelerate PGK1 activity^10^ and therefore considerable ambiguity persists as to whether the protective impact of TZ in both humans^12^ and animals models^11^ is attributable to PGK1 enhancement. Given our findings that boosting PGK1 expression in neurons led to a rescue of hypometabolic function, we sought to determine whether increasing PGK1 expression *in-vivo* would also provide protection against lesions known to lead to PD-resembling axonal degeneration. Stereotactic unilateral injection of 6-hydroxy-dopamine (6-OHDA) into the medial forebrain bundle in mice leads to progressive loss of striatal dopamine axons that is accompanied by asymmetric sensitivity to dopamine receptor (D2R) agonists. Although the precise mechanism of 6-OHDA-driven axon loss is debated, recent experiments support the idea that degeneration begins in the axonal projections^22^. To test whether increasing PGK1 expression would protect against 6-OHDA lesioning, 30 days prior to 6-OHDA injection, an AAV expressing PGK1-mRuby was injected into the Substantia Nigra compacta (SNc) on the same side of the brain that will later receive the 6-OHDA injection (Fig. 3A). In control (PBS injected in SNc, 6-OHDA lesioned) mice, application of the D2R agonist apomorphine led to a robust inducement of whole-body rotations, which was significantly suppressed in animals that received the AAV injection (Fig. 3B). Retrospective histology carried out after the behavioral tests showed that the number of TH (Fig. 3C, D) and DAT (Fig. 3E, F) positive neurons in the SNc was ∼2.5 fold higher in animals that received the AAV harboring PGK1, and that most of these DA neurons also stained positive for tagged PGK1 (Fig. 3C, E). In addition, retrograde tracer injected in the striatum demonstrated considerable protection of intact nigrostriatal neurons compared with controls, with most of the tracer co-localizing with tagged PGK1, (Fig. 3G, H), further suggesting that PGK1 expression protects the axonal integrity of dopaminergic neurons.

**Figure 3.**
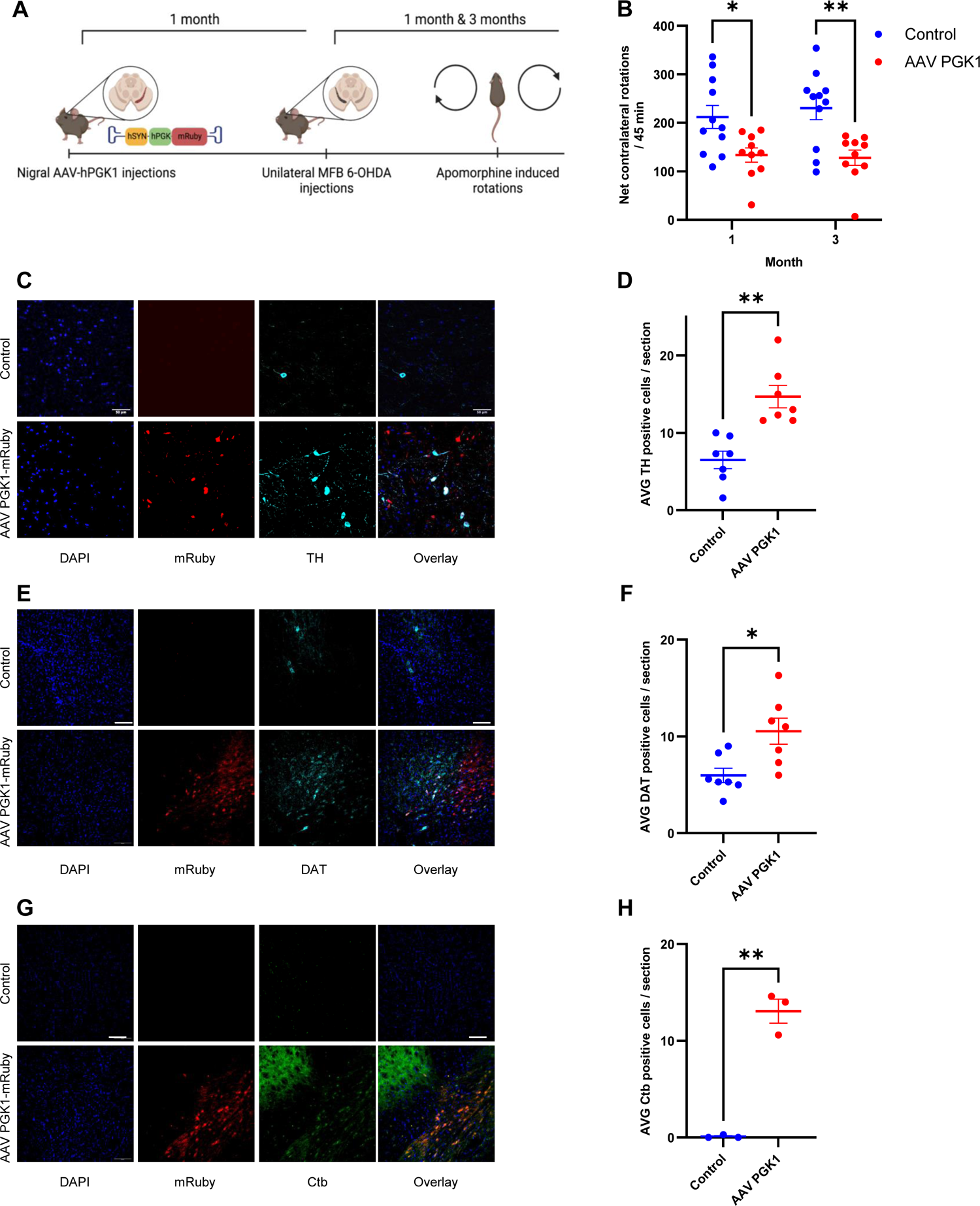
PGK1 protects dopaminergic neurons degeneration *in vivo*. **(A)** An AAV driving hSyn PGK1-mRuby was injected unilaterally in the substantia nigra 30 days before 6-OHDA was injected in the medial forebrain bundle, with control mice only receiving the 6-OHDA injections. At 30 or 90 days after the 6-OHDA injection, both controls and AAV injected mice were subjected to apomoprhine induced rotation tests. (**B**) PGK1-mRuby expression significantly suppressed the apomorphine induced rotations, mean ± SEM, Control N=11, AAV PGK1 N=10 *p < 0.05, **p < 0.01 2-way ANOVA. (**C**) Immunostaining of control and AAV PGK1 mid-brain brain slices against a nuclear marker, DAPI, mRuby and a dopaminergic neuronal marker, TH. Scale bar 50 μm. (**D**) Quantification of the number of TH positive cells showed a significant increase in the amount of alive cells post lesion in case of the PGK1 AAV. Control N=8, AAV PGK1 N=8 **p < 0.01 unpaired t-test. (**E**) Immunostaining of control and AAV PGK1 mid-brain brain slices against a nuclear marker, DAPI, mRuby and a second dopaminergic neuronal marker, DAT. Scale bar 100 μm. (**F**) Quantification of the total number of DAT positive cells showed a significant increase in the amount of alive cells post lesion in case of the PGK1 AAV. Control N=7, AAV PGK1 N=7 *p < 0.05 unpaired t-test. (**G**) Prior to carrying out retrospective immunostaining, the striatums of the animals were injected with a retrograde tracer, cholera toxin subunit b, Scale bar 100 μm. (**H**) Quantification of the number of Ctb positive cells also showed a significant increase in the amount of alive cells post lesion in case of the PGK1 AAV. Control N=3, AAV PGK1 N=3 **p < 0.01 unpaired t-test.

### Terazosin restores synaptic function under hypometabolic conditions

As one of the starting points for implicating a pivotal role for PGK1 in PD was the identification of PGK1 as an off-target binder of TZ, we examined whether TZ mimicked PGK1 expression in our synaptic endurance paradigm. Treating neurons with 10 μM TZ conferred significant resilience to stimulation in low glucose (Fig. 4A, B) albeit to a lesser extent than with PGK1 expression (see discussion). This impact was eliminated in neurons where PGK1 expression was depleted by expression of an shRNA targeting PGK1 (Fig S2D, E). Consistent with *in-vitro* enzymology^10^, a lower dose (1 μM) was ineffective, while a higher dose (100 μM) was either ineffective or likely deleterious, since we found few cells that survived this treatment. TZ’s impact on synaptic metabolism was also independent of the known clinical target of the drug, the α1 adrenergic receptor (α_1_R), as a modified version of TZ that lacks potency for α_1_R, TZ-md^23^, was equally effective at conferring synaptic endurance (Fig S2F, G). Parallel ATP measurements showed that TZ lead to improved nerve terminal ATP production as well. In neurons treated with TZ, resting ATP levels were slightly elevated, ATP depletion during strong stimulation in low glucose was blunted, and the recovery following such a stimulus was accelerated (Fig 4C, D) compared to controls. These data collectively support the concept that PGK1 under low glucose conditions is rate-limiting for ATP production at nerve terminals, and that TZ increases PGK1 activity and thereby ATP production.

**Fig. 4.**
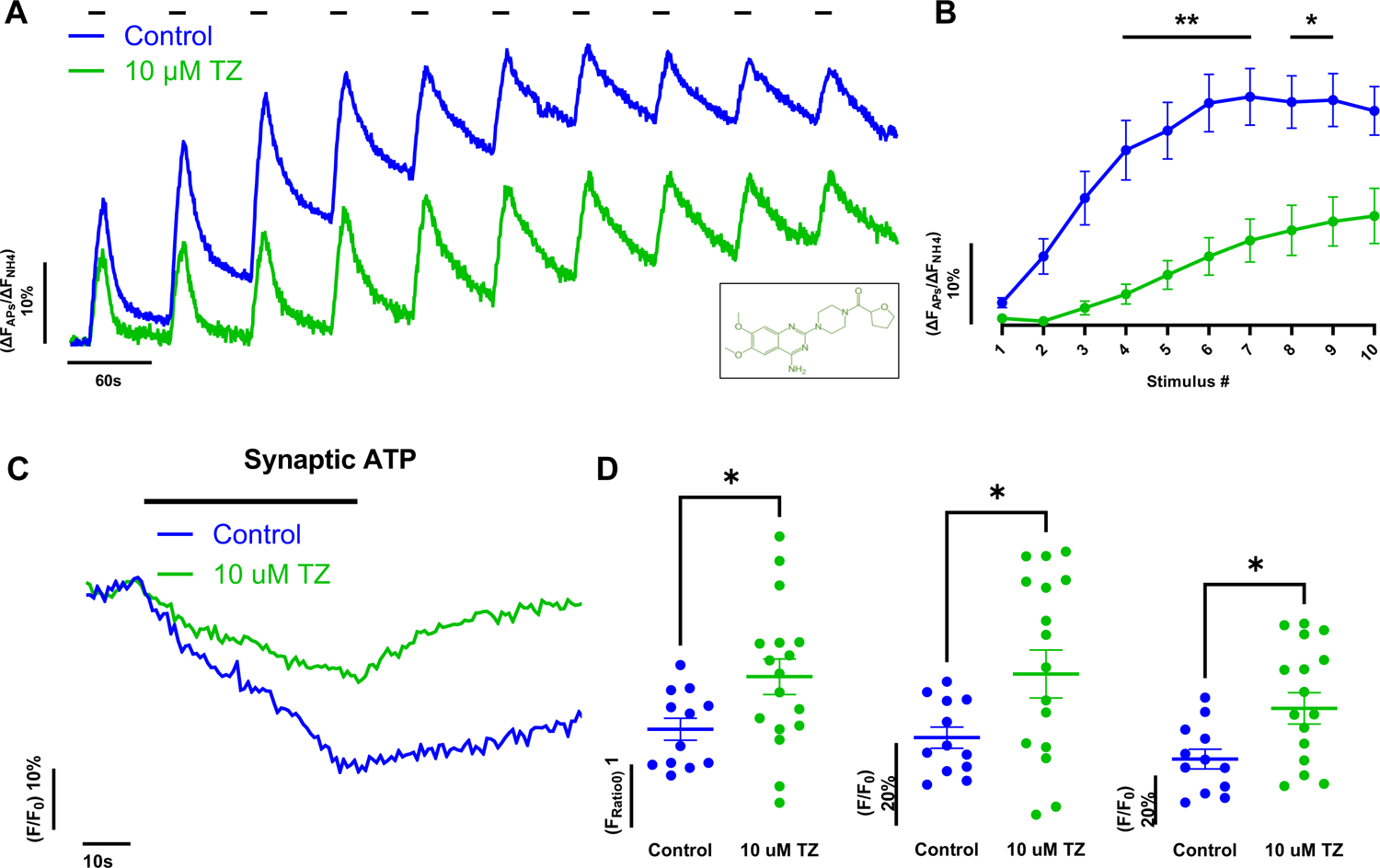
Terazosin confers metabolic synaptic resilience. **(A)** Ensemble average vGlutI-pH traces for control (blue) and Terazosin (TZ) treated (green) primary hippocampal neurons subjected to repeated stimulation in 0.1 mM glucose (TZ chemical structure shown in window). (**B**) The synaptic endurance, measured as the remaining fluorescence 55 sec after each AP bout (mean ± SEM) for the traces in (**A**) show that TZ confers significant resilience. Control N=23, Terazosin N=10 *p < 0.05, **p < 0.01 2-way ANOVA. (**C**) The kinetics of presynaptic ATP measured with synapto-iATPSnFR2-miRFP670nano3, normalized to the pre-stimulus values reveal that TZ incubated cells stimulated with 600 APs in 0.1 mM glucose have significantly smaller activity-induced drops in ATP and more rapid post-stimulus recovery as well as higher starting ATP. (**D**) Comparison of the absolute nerve terminal ATP values pre-stimulus (left) at the end of the stimulus (middle) and 35 s post-stimulus (right) in control and TZ treated neurons mean ± SEM indicated, control N=12, TZ N=17. *p < 0.05, **p < 0.01 unpaired t-test.

### PARK7 (DJ-1) interacts with PGK1 and is necessary for PGK1 mediated synaptic resilience

An unbiased chemical screen recently led to the identification of an interaction between PGK1 and DJ-1^19^, the product of the PARK7 gene, a genetic driver of familial PD^24,25^. DJ-1 is a chaperone with strong structurally similarity to HSP31^26^ but the precise clientele that it supports related to PD is poorly understood^27^. To investigate whether DJ-1 has a functional impact on PGK1 function, we probed PGK1 rescue of hypometabolic synaptic function, but in absence of DJ-1. Although shRNA-mediated loss of DJ-1 led to a very similar gradual slowing of SV recycling in low glucose under repeated stimulation to that seen in control neurons (Fig, S3A, B, G) over-expression of PGK1 or incubation with TZ in absence of DJ-1 completely failed to restore synaptic endurance under low glucose conditions (Fig. 5A, B, C, D). The need for DJ-1 to confer PGK1-dependent metabolic resilience strongly suggests, that loss of DJ-1 might itself impact synaptic ATP dynamics during activity. To test this idea, we carried out measurements of presynaptic ATP dynamics in DJ-1 KD neurons using synapto-iATPSnFR2.0-miRFP670nano3. These experiments revealed a striking inability of DJ-1 KD nerve terminals to sustain ATP production during stimulation under hypometabolic conditions (Fig. 5E). During prolonged stimulation in low glucose, ATP levels dropped by 60%, more than double that in control neurons and exhibited no recovery thereafter (Fig. 5F). Consistent with the inability of PGK1 activity enhancement to rescue hypometabolic synapse function, TZ application showed no acceleration of ATP production in DJ-1 KD neurons following a prolonged burst of activity (Fig. 5G, H). The impairment in ATP production resulting from loss of DJ-1, was likely due to a loss of glycolytic function and not mitochondrial ATP production as a slowing of synaptic vesicle recycling in low glucose and during mild stimulation was only apparent when we additionally blocked mitochondrial ATP production with the F_1_-F_0_ ATPase inhibitor, oligomycin (Fig. S3C, D, F). Furthermore, when glucose was substituted with a mixture of lactate and pyruvate, SV recycling kinetics in DJ-1 KD neurons was indistinguishable from control (Fig. S3E, F). As expected from the impact on ATP production due to the loss of DJ-1, SV recycling in DJ-1 KD neurons was dramatically slowed (Fig. S4A) following intense stimulation. This slowing of SV recycling was reversed upon re-expression of an shRNA-insensitive DJ-1 variant (Fig. S4A, B) but could not be reversed by PGK1 expression or incubation with TZ (Fig. S4C).

**Fig. 5.**
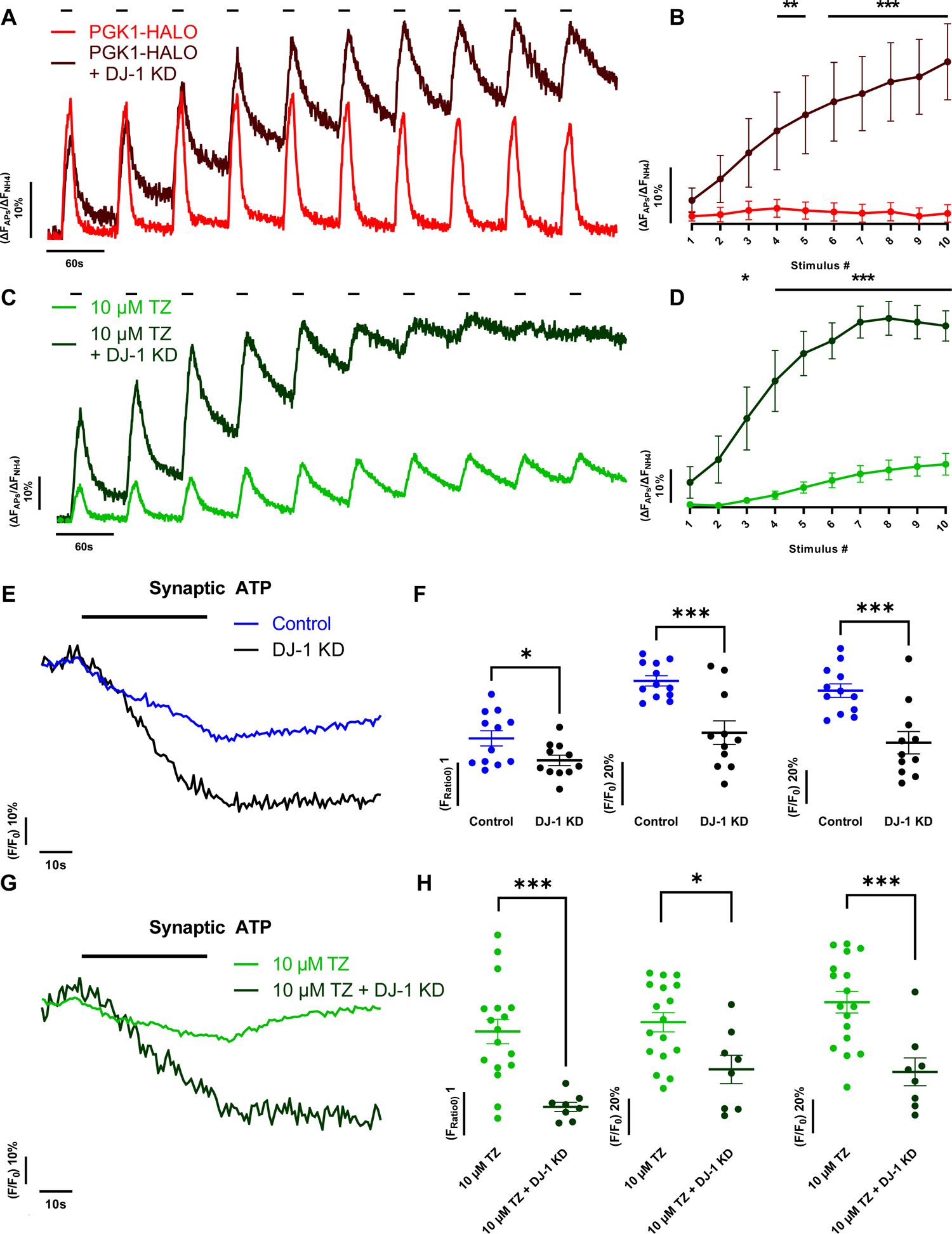
PARK7/DJ-1 is necessary for PGK1 driven hypometabolic resilience. (**A**) Ensemble vGlutI-pH traces in neurons, expressing PGK1-HALO (red) or PGK1-HALO with DJ-1 KD (black) and (**C**) TZ treated neurons (green) versus TZ treated in DJ-1 KD (black) subjected to repeated electrical stimulation (**B**) Synaptic endurance measured vGlutI-pH fluorescence 55 s after each stimulus bout for the traces in (**A**) mean ± SEM, PGK1-HALO N=12, PGK1-HALO + DJ-1 KD N=7 **p < 0.01, ***p < 0.001 2-way ANOVA and (**D**) mean ± SEM, Terazosin N=10, Terazosin + DJ-1 KD N=10 \**p* < 0.05, \*\*\**p* < 0.001 mixed-effects (**E**) The kinetics of presynaptic ATP normalized to the pre-stimulus values when neurons were stimulated with 600 APs at 10 Hz in 0.1 mM glucose in absence of DJ-1 reveal a significantly larger activity-induced drop in ATP and defective post-stimulus recovery. (**F**) Comparison absolute nerve terminal ATP values pre-stimulus (left) at the end of the stimulus (middle) and 35 s post-stimulus (right) in control and DJ-1 KD neurons mean ± SEM indicated, control N=12, DJ-1 KD N=11. *p < 0.05, ***p < 0.001 unpaired t-test. (**G**) The kinetics of presynaptic ATP normalized to the pre-stimulus values when neurons were stimulated with 600 APs at 10 Hz in 0.1 mM glucose in TZ and DJ-1 KD reveal an inability of TZ to accelerate ATP kinetics in absence of DJ-1. (**H**) Comparison absolute nerve terminal ATP values pre-stimulus (left) at the end of the stimulus (middle) and 35 s post-stimulus (right) in TZ and TZ with DJ-1 KD neurons mean ± SEM indicated, TZ N=17, TZ DJ-1 KD N=8. *p < 0.05, ***p < 0.001 unpaired t-test.

These findings suggest that DJ-1 controls PGK1 activity. This could be direct, i.e. that PGK1 is itself a bona fide client of DJ-1, or indirect, perhaps by a previously proposed DJ-1 mediated prevention of 1,3 bisphophoglycerate (1,3BP) driven post translational modifications of numerous proteins, including enzymes that participate in glycolysis ^28^. To investigate a potential interaction between DJ-1 and PGK1 we used micro-scale thermophoresis of recombinant purified DJ-1 and PGK1 that showed an *in-vitro* interaction (∼ apparent K_d_ ∼ 15 μM) (Fig. 6A), in agreement with the findings in the unbiased chemical screen^19^. Furthermore, in axons, DJ-1, is co-localized with PGK1 (Fig. 6B) which supports the plausibility that DJ-1 and PGK1 might interact in the cellular milieu. One possible explanation of the PGK1 activity dependence on DJ-1 is that loss of DJ-1 might in turn lead to a depletion of PGK1, as might be expected if DJ-1 acts as a chaperone to prevent PGK1 degradation. To examine this, we carried out quantitative immunostaining against nerve terminal PGK1 in DJ-1 KD neurons. These experiments demonstrated, that contrary to what would be predicted from a simple proteostasis maintenance role of DJ-1, PGK1 levels increased ∼ 2-fold when DJ-1 is depleted (Fig. 6C). This upregulation of PGK-1 abundance suggests that a feedback loop sensing PGK1 activity may be responding to the lack of DJ-1. Remarkably, when we depleted PGK1 in neurons (via shRNA-mediated KD) quantitative immunostaining against nerve terminal DJ-1, shows that its abundance is upregulated by ∼2-fold under these conditions (Fig. 6D). This reciprocal relationship strongly implies that the measured *in-vitro* interaction likely reflects a close functional relationship between these two proteins. The complete inability of PGK1 to support hypometabolic synaptic function, when DJ-1 is silenced (Fig. 5A, B, C, D) as well as the exaggerated ATP depletion during activity strongly implies that PGK1 may be a *bona fide* client of DJ-1 chaperone activity. These data establish that DJ-1 is necessary for PGK1 function when fuel becomes limiting.

**Fig. 6.**
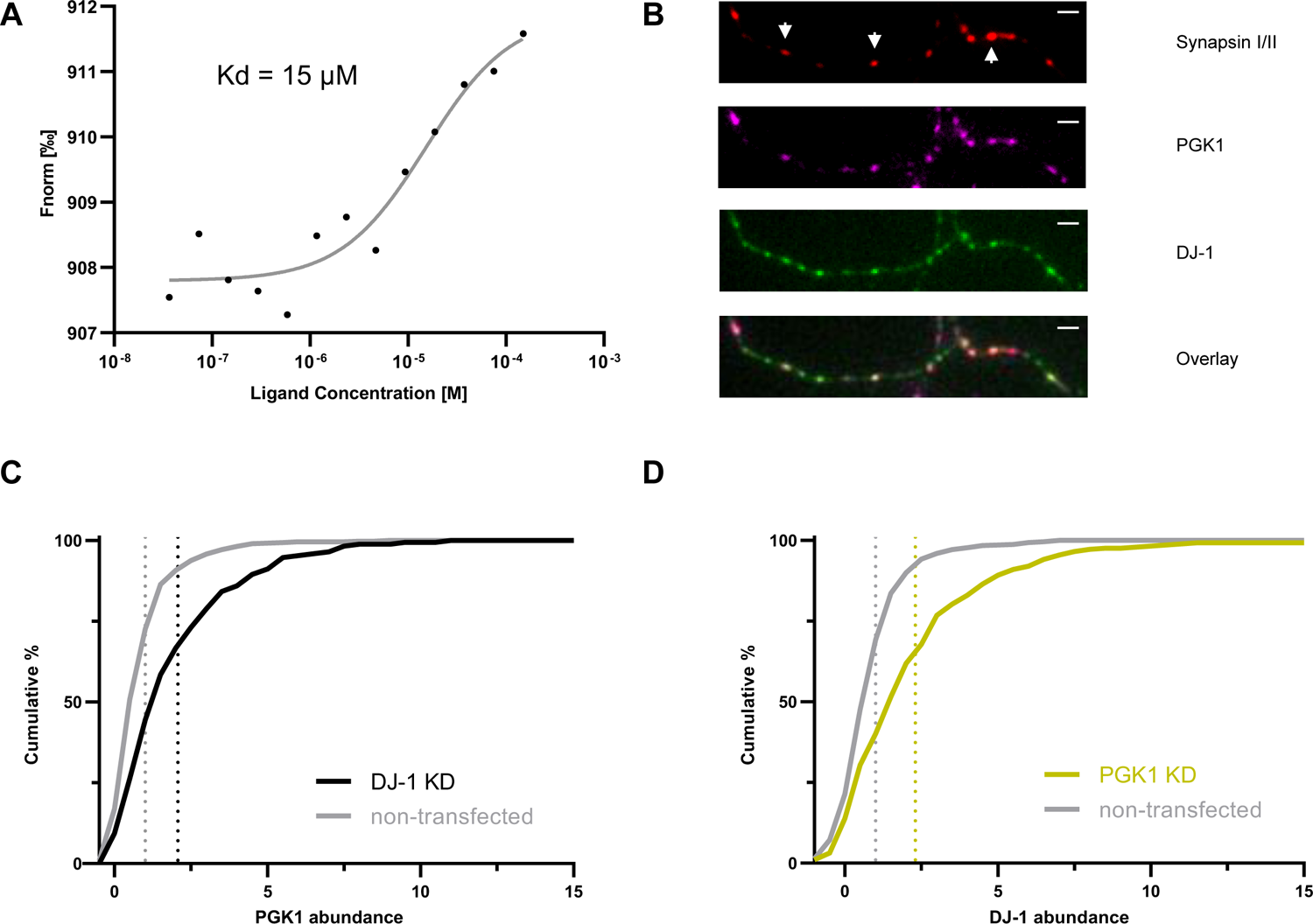
DJ-1 interacts with PGK1 while their levels are co-regulated in nerve terminals. **(A)** Microscale thermophoresis identified an *in-vitro* interaction between PGK1 and DJ-1 with a low micromolar affinity (∼ 15 μM). (**B**) Immunostaining against PGK1 (magenta), synapsin (red) and DJ-1 (green) shows that DJ-1 and PGK1 are both present in nerve terminals (white arrows), white bar 2 μm. (**C**) Cumulative histogram of presynaptic PGK1 fluorescence immunostaining intensity in control (grey) and DJ-1 KD nerve terminals (black), DJ-1 KD N=171, non-transfected N=500 (**D**) DJ-1 fluorescence immunostaining in control (grey) and PGK1 KD nerve terminals (yellow), PGK1 KD N=290, control N=500). Distributions in both KD are significantly different then their respective controls ***p < 0.001 Kolmogorov-Smirnoff.

### PGK1 expression restores PARK20 synaptic dysfunction

Our data suggest an important role for PGK1 in synaptic function as well as a protective role in neurodegeneration. To directly examine the ability of PGK1-HALO to rescue an alternate genetically driven PD phenotype, we used primary neuronal cultures from PARK20 mice, which harbor the R258Q mutation in Synaptojanin I, and are characterized by synaptic abnormalities and impaired SV recycling^29^. While expression of PGK1-HALO in WT neurons had no impact on SV recycling kinetics (Fig. 7A), it restored the slow SV recycling kinetics in PARK20 nerve terminals (Fig. 7B, C). These experiments suggest that the PD driving mutation in Synaptojanin I either creates a metabolic burden, or impairs ATP production in a fashion that can be bypassed by PGK1 overexpression.

**Fig. 7.**
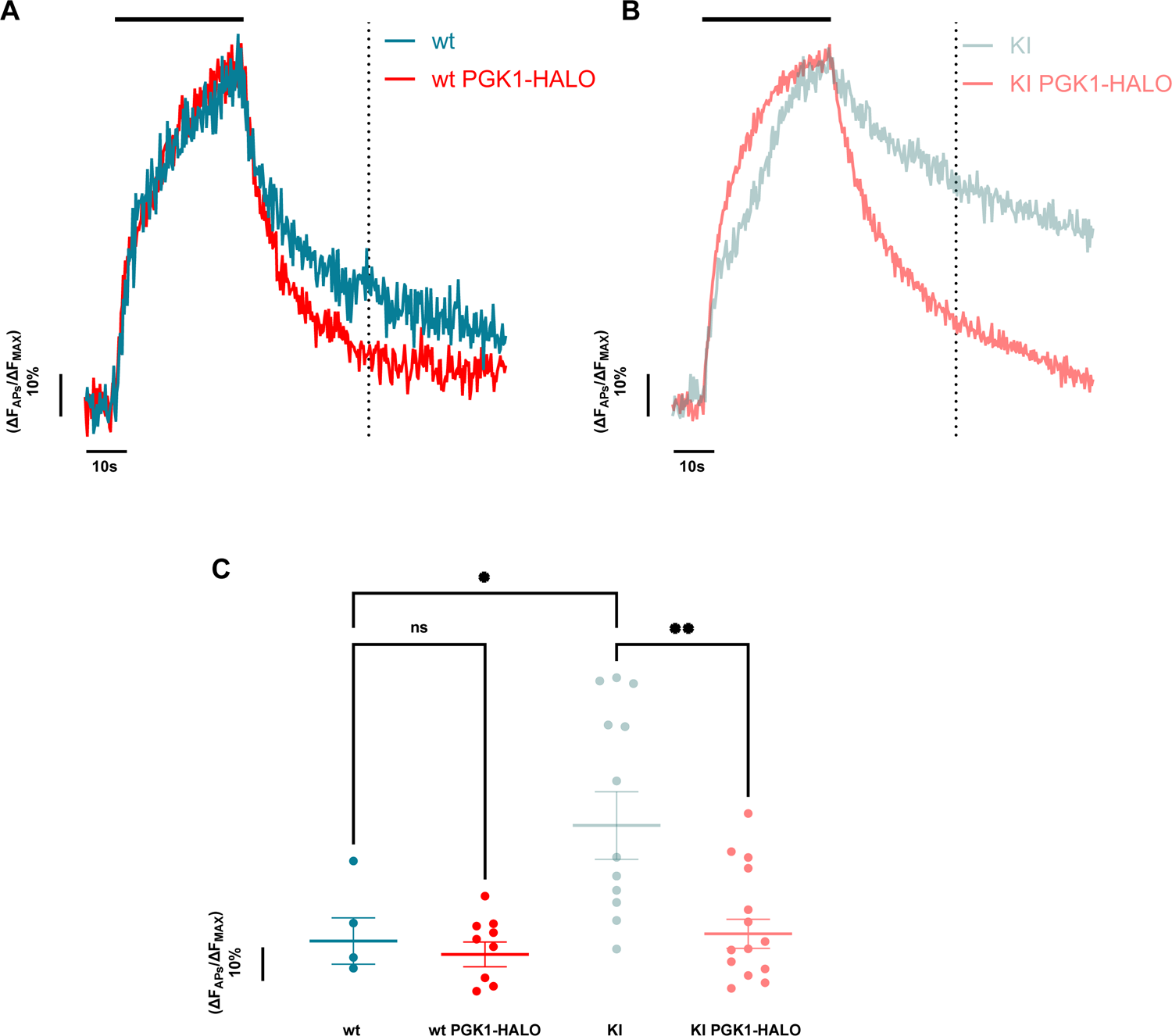
PGK1 restores synaptic function in PARK20 neurons (SynjI R258Q mutation) **(A)** Ensemble average vGlutI-pH traces in primary wt and (**B**) SynjI R258Q KI mouse cortical neurons stimulated with 600 APs (10 Hz, indicated by the black bar) with PGK1-HALO expression (red) or without (teal). (**C**) Quantification of the remaining fluorescence of the vGlut-pH traces 60 s post stimulation (indicated by the dotted line) mean ± SEM, wt N=6, wt PGK1-HALO N=10, KI N=12, KI PGK1-HALO N=16 *p < 0.05, **p < 0.01 1-way ANOVA.

## Discussion

The metabolic vulnerability of the human brain is evident in its general intolerance to reduction in fuel availability. This likely arises for two reasons. The first is that the brain consumes far more of its share of fuel per weight than any other organ in the body. The second is that synaptic transmission, the central control point in information flow in the brain, is one of the loci of this vulnerability. It has long been suspected that an important driver of PD results from a combination of a general age-related decline in the ability to adequately fuel the brain, the outsized vulnerability of highly arborized and energetically demanding SNc DA neurons, and genetic lesions that impair either the efficacy of ATP production or its use within the brain. To date, human genetic studies have led to the identification of at least 23 drivers of familial PD, the PARK genes^30^. Although two of the PARK genes, PARK2 (PARKIN) and PARK6 (PINK1) are potentially directly linked to bioenergetics as they participate in mitochondrial quality control^5^, the connection between other PARK genes and bioenergetics is less clear. The discovery that TZ is both therapeutically protective in humans^12,17^ as well as numerous animals models^11^ provided some of the most compelling evidence that bioenergetic dysfunction might be central to PD. One challenge in interpreting these previous findings is knowing whether or not the protective impact of TZ was really attributable to and enhancement of PGK1 activity, since the *in-vitro* data only showed a very modest impact of TZ on PGK-1 function^10^. Our data speak directly to this and demonstrate both that enhancing PGK1 activity protects striatal DA axons in vivo, and dramatically impacts the kinetics of subcellular ATP production.

The biochemistry of glycolysis has been established for many decades and numerous regulatory feedback loops and control points are well understood. The serendipitous discovery that TZ can accelerate PGK1 activity, and that TZ is protective against PD in humans and numerous experimental PD models implies that PGK1 is a rate-limiting enzyme in glycolysis, at least in neurons. In this study we exploited the metabolic sensitivity of presynaptic function in primary neurons, to establish a sensitized real-time assay of synaptic endurance. The logic of this approach is that increasing the abundance of the rate-limiting enzyme should lead to improved function in a sensitized assay thus allowing us to test the prediction arising from the TZ discovery. Although negative results from our low-throughput expression screen are not easily interpreted (as is true for most genetic screens), a positive result leads to the unambiguous conclusion that the rate-limiting step has been identified. The direct test that upregulation of PGK1 abundance improves glycolysis was made possible with the recent development of a next-generation ATP sensor^21^, which revealed that following acute ATP depletion driven by intense activity, the restoration of ATP levels in low glucose conditions was dramatically accelerated by PGK1 expression. Our experiments with TZ were in broad agreement with the original data describing TZ’s neuroprotective effects^11^. Two previous studies showed that PGK1 overexpression is neuroprotective. In a mouse model of spinal muscular atrophy (SMA)^18^, Boyd and colleagues found a strong correlation between PGK1 expression and the resilience to spinal cord motor neuron degeneration and that overexpression of PGK1 was strongly protective in a zebrafish model of this genetic disease. Similarly, PGK1 expression in drosophila neurons, but not muscle, protected flies from rotenone-induced motor neuron dysfunction^11^. Here, we show that PGK1 expression is neuroprotective in the specific neurons that are vulnerable in PD (Fig. 3) using an established model (6-OHDA) that drives mammalian striatal axon degeneration.

Our data also reveal an unexpected link between PGK1 and PD, as the ability to accelerate the kinetics of ATP production by enhancing PGK1 activity was completely dependent upon DJ-1, a known genetic driver of PD in humans. Our data are consistent with the idea that there is a functional partnership between DJ-1 and PGK1 whereby DJ-1 directly modulates PGK1 activity. This interpretation is supported both by our *in-vitro* binding data and our finding that when either PGK1 or DJ-1 levels are depleted it leads to a corresponding increase in the other protein (Fig. 6). This apparent cross-regulation of these two proteins suggests that rather than acting as a classical chaperone to simply restore proper client folding and preventing protein degradation, it acts instead as a chaperone to sustain enzyme activity.

A recent analysis demonstrated that loss of DJ-1 led to an increase in a novel form of post-translational lysine glyceration throughout the proteome in DJ-1 knock cells^28^, a modification that was previously shown to be driven non-enzymatically by the substrate for PGK1 in glycolysis (1,3 BP)^31^. This analysis also showed that although DJ-1 could prevent lysine glyceration from accumulating in the presence of 1,3 BP *in-vitro*, it could not remove this modification once it had formed^28^. These data imply that, contrary to previous views that DJ-1 is a glycase^32^, DJ-1 acts instead to prevent the formation of this post-translational modification and does not catalyze its removal. At present it is unclear what the potential consequences of lysine glyceration are on proteins that accumulate this modification, several of which were glycolytic proteins themselves. Our data that ATP levels become severely depleted at nerve terminals during activity in the absence of DJ-1 are also consistent with the interpretation that these lysine glycations on glycolytic enzymes would impair their activity, in turn impairing ATP production.

Previous studies showed that TZ was remarkably protective in several genetic models of PD^11^, including PARK4, PARK6 and PARK8. However, the interpretation of this data hinged upon the previous identification of the off-target binding of TZ to PGK1. The *in-vitro* analysis in that work reported that TZ could accelerate PGK1 activity, but only by ∼ 5%^10^, casting doubt on whether the *in-vivo* impact of TZ was mediated by PGK1. Our studies unambiguously show that enhancing PGK1 activity is highly protective and complements previous findings by additionally demonstrating that specific synaptic deficits driven by PARK20/Synaptojanin I R258Q can also be reversed by PGK1 expression. These data all imply that a common underlying link between the many disparate genetic drivers of PD is likely bioenergetic in origin. We anticipate that certain mutations compromise ATP production, either for example by impairing mitochondrial function as is implied in PARK2/PARK6 or impairing glycolysis as seems to be occurring with PARK7/DJ-1 and potentially PARK12. Other drivers of PD may result in creating undue metabolic burdens which in turn compromise axon and synapse function, as may be the case for PARK20/Synaptojanin-1. These results establish PGK1 as a crucial leverage point in synaptic function and PD-related dysfunction and demonstrates that neuronal glycolysis is an essential pathway that fuels synaptic transmission. Therefore, the mechanisms that control PGK1 abundance, activity and localization, which are not well understood, are of significant therapeutic potential. In humans, PGK1 mutations present usually in three broad clinical deficits characterized by symptoms associated with anemia, myopathy and neurological dysfunction^33^. This strongly suggests that different cells and tissues are differentially vulnerable to different mutations, implying that their local regulation may differ. Further investigation into how local abundance and activity is controlled in different tissues should prove insightful. Our study significantly strengthens the case that PGK1 biology is of direct therapeutic interest for PD and neurodegenerative diseases in general. A detailed understanding of the molecular mechanism underlying PGK1 activation will have the potential to open new therapeutic routes for this disease.

## Acknowledgments

We would like to thank members of the Ryan lab for helpful discussions and Luke Lavis for the gift of JF dyes. This research was supported in part by NIH grants (NS036942 & NS11739) to TAR and in part by Aligning Science Across Parkinson’s ASAP-000580 and ASAP-020608 through the Michael J. Fox Foundation for Parkinson’s Research (MJFF). For the purpose of open access, the authors have applied a CC BY public copyright license to all Author Accepted Manuscripts arising from this submission.

## Materials and Methods

### Reagents

Chemical reagents were purchased from Millipore Sigma. Drugs were purchased from Tocris Bioscience. Antibodies used are a-PGK1 abcam ab199438, a-DJ-1 Cell Signaling 5933 (RRID AB_11179085) and Santa Cruz sc-55572(RRID AB_831639), a-synapsin I/II Synaptic Systems 106 004 (RRID AB_1106784), a-GAPDH Cell Signaling 5174 (RRID AB_10622025), a-Tubulin beta III R&D Systems MAB1195 (RRID AB_357520), a-GFP Thermo Fisher A10262 (RRID AB_2534023), a-TH Millipore AB1542 (RRID AB_90755), a-DAT Proteintech 22524-1-AP (RRID AB_2879116). Alexa Fluor-conjugated fluorescent secondary antibodies were obtained from Life Technologies. Terazosin was purchased from Tocris Bioscience (1506).

### Animals

All animal-related experiments were performed in accordance with protocols approved by the Weill Cornell Medicine Institutional Animal Care and Use Committee. Wild-type rats were of the Sprague-Dawley strain (Charles River Laboratories strain code: 400, RRID: RGD_734476). The prak20/Synaptojanin R258Q KI mice^29^ were maintained in the De Camilli lab. For behavioral experiments, male C57BL/6J mice (Jax.org; strain #:000664) were housed two to five per cage and kept at 22°C on a reverse 11 am-light/11 pm-dark cycle, with standard mouse chow and water provided ad libitum throughout the duration of the study.

### Plasmids

The following previously published DNA constructs were used: vGLUT1-pHluorin^34^, DJ-1 (a gift from Mark Cookson, Addgene plasmid 29347), synapto-iATPSnFR2-miRFP670nano3. For this study these new plasmids were generated: mTagBFP2-N1, PLKO mTagBFP2, PGK1 KA, DJ-1 KA, PGK1-HALO, PGK1-mRuby.

### Primary neuronal culture

Primary hippocampal CA1 to CA3 neurons were isolated from day 1 rat pups of mixed gender and plated on poly-l-ornithine–coated coverslips as previously described^35^ in minimum essential medium (Thermo Fisher 51200038) supplemented with 0.5% glucose, 0.1 g/L bovine transferrin, 0.24 g/L insulin, 1% Glutamax (Thermo Fisher 35050061), 5% fetal heat inactivated FBS (Atlanta Biologicals S11510) and 2% homemade N-21^36^ in a humidified incubator at 37°C 5% CO2. After 2 days the media was changed to culture media containing in addition 4 μM cytosine b-d-arabinofuranoside. On day 6, cells were transfected using calcium phosphate as previously described^37^.

### Live-cell imaging

Live-cell imaging was performed using a custom-built laser-illuminated epifluorescence microscope with an Andor iXon+ (model no. DU-897E-BV) back-illuminated electron-multiplying charge-coupled device camera. Transistor– transistor logic (TTL)-controlled Coherent OBIS 488-, 561- and 637-nm lasers were used for illumination. Images were acquired through a 40× 1.3 numerical aperture (NA) Fluar Zeiss objective. Experiments were performed at a clamped temperature of 37°C using a custom-built objective heater under feedback control. Action potentials were evoked by passing 1-ms current pulses, yielding fields of approximately 10 V cm−1 via platinum-iridium electrodes. Neurons were continuously perfused at 0.1 ml/min with a Tyrode’s solution containing 5 mM glucose, 50 mM Hepes, 119 mM NaCl, 2.5 mM KCl, 2 mM CaCl_2_, 2 mM MgCl^2^, 0.01 mM 6-cyano-7-nitroquinoxaline-2,3-dione, 0.05 mM d,l-2-amino-5-phospho-novaleric acid pHed at 7.4 for high glucose experiments. For low glucose, the tyrodes buffer used contained 0.1 mM glucose, 54.9 mM Hepes, 119 mM NaCl, 2.5 mM KCl, 2 mM CaCl_2_, 2 mM MgCl^2^, 0.01 mM 6-cyano-7-nitroquinoxaline-2,3-dione, 0.05 mM d,l-2-amino-5-phospho-novaleric acid (pH 7.4). When Oligomycin was used it was added at 2 uM (Millipore Sigma O4876). For vGlutI-pH normalization, cells were perfused with Tyrodes containing 50 mM NH_4_Cl pHed at 7.4.

### Suppressor screen

Primary hippocampal neurons co-transfected with vGlutI-pH and a plasmid encoding one of the following glycolytic enzyems: Hexokinase 1, Glyceraldehyde phosphate dehydrogenase, PGK1, Alodolase A, Aldolase C or Pyruvate kinases M1. Neurons were incubated in imaging media containing 0.1 mM Glucose for 5 minutes after mounting into the stimulus chamber. Neurons were then challenged with ten bouts of AP firing (100 APs at 10 Hz) delivered every minute. Cells were finally perfused with imaging media containing 50 mM NH_4_Cl.

### vGlutI-pH analysis

For vGlutI-pH experiments, nerve terminals responding to the first train of action potentials were selected and background subtracted. In case of the repeated stimulation assay, the traces are reported as ΔF by subtracting the initial fluorescence signal before stimulation and normalized to total sensor fluorescence, revealed by the 50 mM NH_4_Cl Tyrodes solution, yielding ΔF_APs_/ΔF_NH4_. For single train experiments, the traces are reported ΔF_APs_/ΔF_MAX_, by normalizing to the fluorescence peak during the action potential train. The remaining fluorescence measurements are calculated at the specified time point for each data-set. All graphs were made with GraphPad Prism 10.0.2 and sketches with BioRender.

### ATPsnfr analysis

For synapto-iATPSnFR2-miRFP670nano3experiments, nerve terminals were selected in the miRFP670nano3 channels, blind to the ATPsnfr channel and then background subtracted. The traces are reported as iATPsnfr2 to miRFP670nano3 ratio, F_ratio_. Cell data was filtered by removing any cells that exhibited no change to activity.

### Immunocytochemistry

For immunostaining experiments, we used a modified culture protocol to seed cells in a lower density. These cells were instead plated on poly-D-lysine treated coverslips in absence of cylinders and transfected using a Lipofectamine 2000 reagent, as previously described^38^. Cells on a coverslip were fixed using 4% PFA, treated with 50 mM NH_4_Cl and then permeabilized using 0.1% Triton X-100. Cells were blocked using a 5% BSA buffer and the antibodies were incubated in the same buffer. Finally, cells were mounted on glass slides using Prolong Diamond (Thermo Fisher P36970). For shRNA quantification, regions of interested including the cell somas were selected. For presynaptic analysis, regions of interested that were synapsin I/II positive/negative were selected blindly to the other channels.

### AAV craniotomy and *in-vivo* PD model

Stereotactic surgical procedures were performed on 8 - 12-week old mice, weighing 20 - 25g, under a mixture of ketamine/xylazine anesthesia. Ketamine (Butler Animal Health Supply) and xylazine (Lloyd Laboratories) were administered at concentrations of 110 and 10 mg/kg body weight i.p., respectively. After the induction of anesthesia, the animals were placed into a stereotactic frame (David Kopf Instruments). All AAV infusions were performed using a 10 μL stereotactic syringe attached to a micro-infusion pump (World Precision Instruments) at a rate of 0.1 - 0.4 μL/min. To prevent reflux, after each infusion, the injection needle was left in place for 5 min, withdrawn a short distance (0.3 - 0.5 mm), and then left in the new position for an additional 2 min before removal.

hSyn hPGK1-mRuby AAV or PBS were injected unilaterally into the mice substantia nigra pars compacta (AP −3.0 mm, ML ±1.2 mm, DV −4.5 mm from bregma). All craniotomies were performed at a rate of 0.4 μL /min using a 10 μL WPI syringe with a 33 G needle. A total of 2×10^11^ genomic particles/μL (3 μL in PBS) of vector was injected in test animals. Controls received 3 μL in PBS, which was used as a control to avoid potential confounding effects of a control AAV expressing a marker gene that could induce or potential toxicity and lead to a false conclusion of PGK-mediated neuroprotection. The mice were allowed a 6 – 8 week recovery before being subjected to the behavioral tests during which maximal transgene expression from AAV vectors is achieved. At the conclusion of the experiments, injection site accuracy within the targeted brain region was determined by immunohistochemistry for the specific transgene expressed by the AAV vector, and mice with mistargeted injections were excluded from analysis before their data were unblinded.

To generate 6-OHDA lesioned mice, animals were injected with a total volume of 0.6 μL of 6-OHDA hydrobromide (Sigma-Aldrich) in PBS with 0.1% ascorbate unilaterally into the medial forebrain bundle (MFB) at a concentration of 2.5 mg/mL and an infusion rate of 0.1 μL/min with a 10 μL Hamilton syringe with a 30 G needle. The coordinates for the injection were AP −1.1 mm, ML −1.1 mm, DV −5.0 mm relative to bregma and the dural surface. Before lesion surgery, the norepinephrine reuptake inhibitor desipramine (25 mg/kg, i.p.) was administered via i.p. injection at least 30 min before 6OHDA infusion, to protect neostriatal and cerebellar noradrenergic neurons from the toxin-induced damage. Mice were allowed 4 weeks to recover before being subjected to the apomorphine-induced a behavioral test to estimate the extent of dopamine depletion in the substantia nigra (see following section).

For striatal injections of cholera toxin subunit-B (CTB) Alexa fluor-conjugated 488 (Invitrogen Cat. No C34775), unilateral injections were performed into the ipsilateral dorsal striatum to the 6OHDA lesion side (AP +0.5 mm, ML ±2.3 mm, DV −3.5 mm from bregma) 1-week prior to euthanizing the animals.

### Apomorphine induced rotation assay

All behavioral tests were run during the dark phase of the mouse daily cycle. The experiments were performed by an examiner blinded to treatment groups. To test for 6-OHDA lesion efficiency, rotational behaviors were performed. In brief, mice were placed in a body harness connected to a transducer/swivel in a bowl-shaped testing arena (rotometer). Full 360° clockwise and counter-clockwise rotations were measured over a 45 min period after drug injection for apomorphine-induced rotations. Apomorphine (Sigma-Aldrich) was dissolved in saline solution, 0.9% NaCl, and 0.1% ascorbate, and administered at a dose of 0.25 mg/Kg via s.c. injection.

EthoVision XT14 was used to analyze offline rotational behaviors. The automatic animal detection with deep learning settings was used to track center to-nose and center to-tail points in defined arenas. Video recordings were used to analyze the following locomotion and rotational parameters: (1) distance travelled, (2) activity percentage (%), (3) rotation >90 degrees clockwise, and counterclockwise. Tracking data was graphed with EthoVision analysis tools.

### Immunohistochemistry

On completion of all assessments, mice were deeply anesthetized with sodium pentobarbital (150 mg/kg, i.p.) and transcardially perfused with 4% paraformaldehyde (PFA). Brains were extracted and post-fixed overnight in 4% PFA, cryoprotected in 30% sucrose, and cut into 30 μm sections using a vibratome (Leica Microsystems). Free-floating sections were treated with antibodies to visualize proteins of interest using immunofluorescence labeling, including Tyrosine Hydroxylase (TH) and Dopamine Transporter Antibody (DAT) to quantify the extent of nigral neurodegeneration, and mRuby fluorescence to estimate the transduction efficiency from our AAV vector.

Images were taken by an epi-fluorescent microscope (Olympus BX61 microscope fluorescent microscope fitted with an Olympus DP71 digital camera) or a confocal microscope (Zeiss LSM 900) and analyzed with Image J software (version 1.52p, NIH, US). For quantification of TH+ cells, coronal sections were sampled at intervals of 90 - 120μm for immunostaining. 3 nigral sections per mouse were identified using the mouse brain atlas of Paxinos (2007) and scanned bilaterally using a 20x objective. For each selected section, three randomly chosen regions of interest (ROIs) with fixed areas were selected for quantification. A Macro Plugin was applied to each section obtained, with a Gaussian blur filter threshold set to 30 and size filter to 20 to remove small objects and low signal cells. TH+ neurons were determined only when clear co-localization with DAPI staining was observed. Automatized detection using Image J software in 7 mice per condition (controls vs treated).

### Microscale Thermophoresis

His-tagged SUMO-PGK1 and DJ-1 full length proteins were expressed in BL21 (DE3) *E. coli* cells. 1 L cultures in LB media with 50 μg/mL kanamycin were grown at 37°C to OD 600 of 0.6 before induction with 0.7 mM isopropyl-β-d-1-thiogalactopyranoside (IPTG) followed by incubation at 22°C for 18 hours. Cells were harvested by centrifugation at 4,000 rpm for 20 minutes and frozen at −80°C. Thawed cell pellets were lysed by sonication on ice in 200 mL of lysis buffer (50 mM Tris-HCl pH 8.0, 350 mM NaCl, 1 mM PMSF, and 5 mM β-mercaptoethanol). Lysates were clarified by centrifugation at 18,000 rpm for 45 minutes. Supernatants containing soluble proteins were applied to a 5 mL NiNTA-agarose gravity column (Qiagen) pre-equilibrated with lysis buffer. Bound proteins were washed with 200 mL wash buffer (lysis buffer with 30 mM imidazole) and eluted in 30 mL elution buffer (lysis buffer containing 150 mM NaCl and 250 mM imidazole). The eluted fractions were analyzed by SDS-PAGE gel electrophoresis and protein concentrations were estimated by UV absorption at 280 nm. His-tagged DJ-1 protein fractions were pooled and dialyzed using dialysis tubing with a molecular weight cut off (MWCO) of 3.5 kDa to eliminate imidazole and concentrated using an Amicon Ultra-50 centrifugal filter unit (10 kDa MWCO). His-SUMO-PGK1 was pooled and dialyzed with a molecular weight cut off (MWCO) of 3.5 kDa in 4 L of SUMO cleavage buffer (50 mM Tris-HCl pH 7.5, 150mM NaCl, and 0.2 mM TCEP) at 4°C for 24 hours. SUMO protease was added and the mixture was rocked slowly at 4°C for 24 h. Cleavage efficiency was monitored at various time points using SDS-PAGE. The fully cleaved mixture was loaded onto a 5 mL NiNTA-agarose column equilibrated with binding buffer without imidazole (50 mM Tris-HCl pH 7.5, 150mM NaCl, 0.2 mM TCEP) and eluted as described above. The flow-through containing cleaved PGK1 was collected. Fractions containing untagged PGK1 protein were concentrated using an Amicon Ultra-50 centrifugal filter unit (30 kDa MWCO).

Purified His-tagged DJ-1 and untagged PGK1 proteins were subjected to further purification using gel filtration with a Superdex200 Increase 10/300 GL size-exclusion column (GE Healthcare) on an AKTA FPLC system (Cytiva). The column was equilibrated and samples were run in buffer (20 mM sodium phosphate, pH 7.4, 0.2 mM TCEP) at a flow rate of 0.4 mL/min at 4°C. Both proteins eluted at the expected volume based on their molecular weights. Peak fractions were collected and analyzed by Coomassie-stained SDS-PAGE. Protein containing fractions were pooled prior to subsequent microscale thermophoresis (MST) analysis. Protein structural integrity was verified using circular dichroism.

To assess the interaction of PGK1 with DJ-1, His-DJ-1 was first labeled with RED-tris-NTA by mixing 100 μL of 200 nM His-DJ-1 with 100 μL of 100 nM RED-tris-NTA in PBS-T buffer and incubating for 30 minutes at room temperature in the dark. PGK1 was prepared in PBS-T buffer at 152 μM concentration. This stock solution was used for a 16-step 1:1 serial dilution in PBS-T buffer with 10 μL final volume in each sample, followed by addition of 10 μL of 100 nM RED-tris-NTA labeled DJ-1 to each sample. After mixing by pipetting up and down, reactions were incubated at room temperature, in the dark, for 30 min and centrifuged at 14,000 rpm for 10 min at 4°C before loading into Monolith NT.115 premium capillaries (NanoTemper). Samples were loaded into the Monolith NT.115 device (NanoTemper) and MST was measured using 40% (RED) LED and medium MST power. Data were processed using the manufactuer’s software (MO.affinity 2.3, NanoTemper) following published protocols^39^. Control experiments included measurements of the binding of the RED-tris-NTA dye to His-DJ-1 and of a dye-labeled AptamerCy5 to AMP using a manufacturer kit (Monolith NT.115 Control Kit RED, NanoTemper).

---

**Fig. S1.**
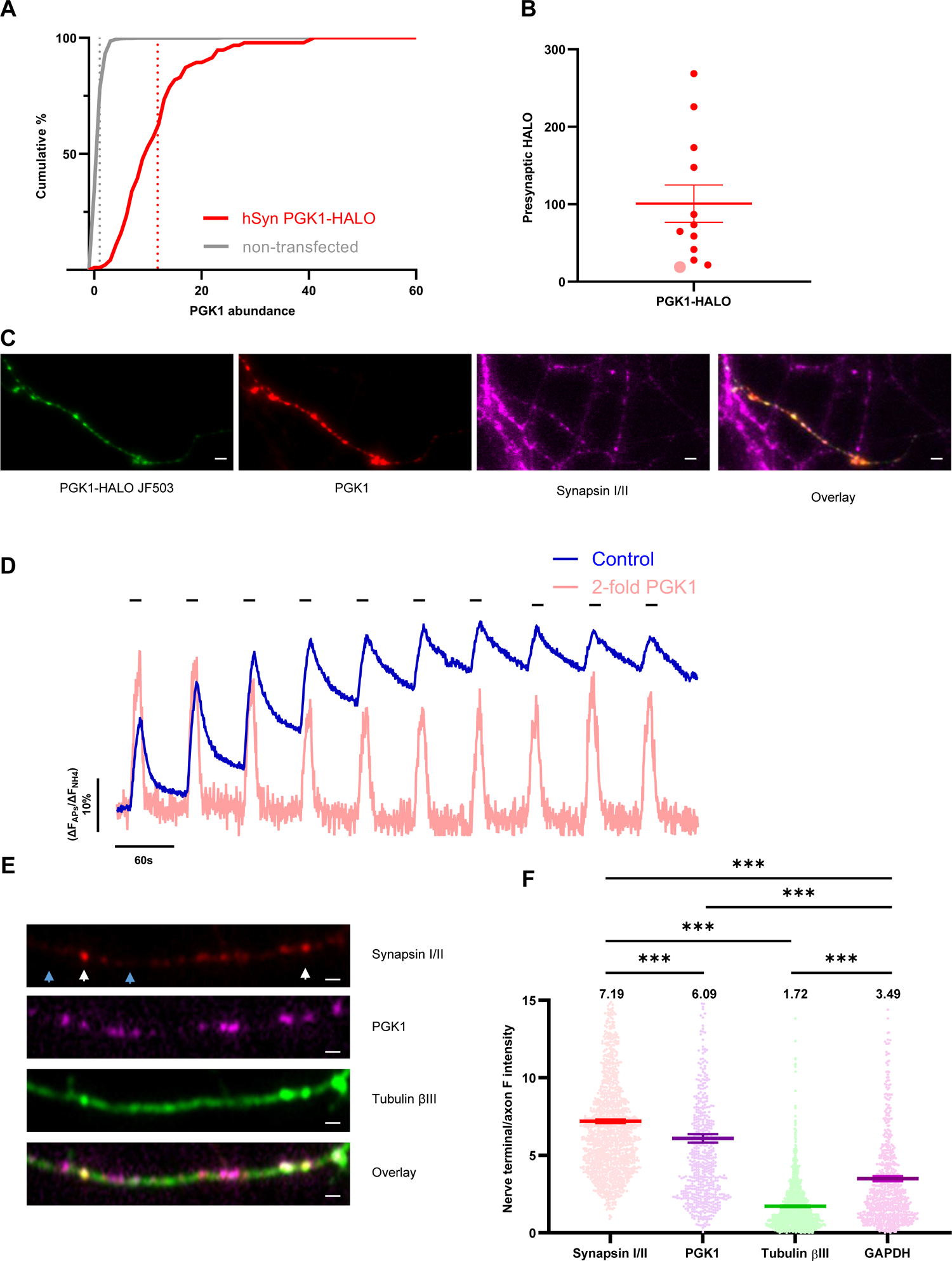
Synaptic PGK1 quantification. **(A)** Synaptic PGK1 was quantified in cells transfected with PGK1-HALO (red) compared to non-transfected cells (grey) at synapsin positive puncta. The construct leads to significant PGK1 overexpression as indicated by the cumulative histograms. PGK1-HALO N=93, non-transfected N=950 puncta \*\*\**p* < 0.001 Kolmogorov-Smirnof. (**B**) The average PGK1-HALO intensity based on JF dye labeling across many experiments, N=12. (**C**) Example of the PGK1-HALO transfected cells labeled with JF503 (green), immunostained against PGK1 (red) and synI/II (magenta). Scale bar is 6 μm. (**D**) vGlutI-pH trace of the lowest PGK1-HALO expressing amount (light red, PGK1 2-fold overexpression, noted as lighter red large point Fig. 2E and S1B) compared to average control (blue). (**E**) Immunostaining against PGK1 (red), a nerve terminal marker synI/II (magenta) and an axonal protein tubulin βΙΙΙ (green). Scale bar is 2 μm. Synaptic (synapsin I/II positive, white arrows) to axonal (synapsin I/II negative, blue arrows) fluorescence intensity ratio is quantified in (**F**) and shows significant enrichment of PGK1 in nerve terminals. mean ± SEM, syn I/II N=1370, Tubulin βIII N=1370, PGK1 N=600, GAPDH N-770 \*\*\**p* < 0.001 1-way ANOVA.

**Fig. S2.**
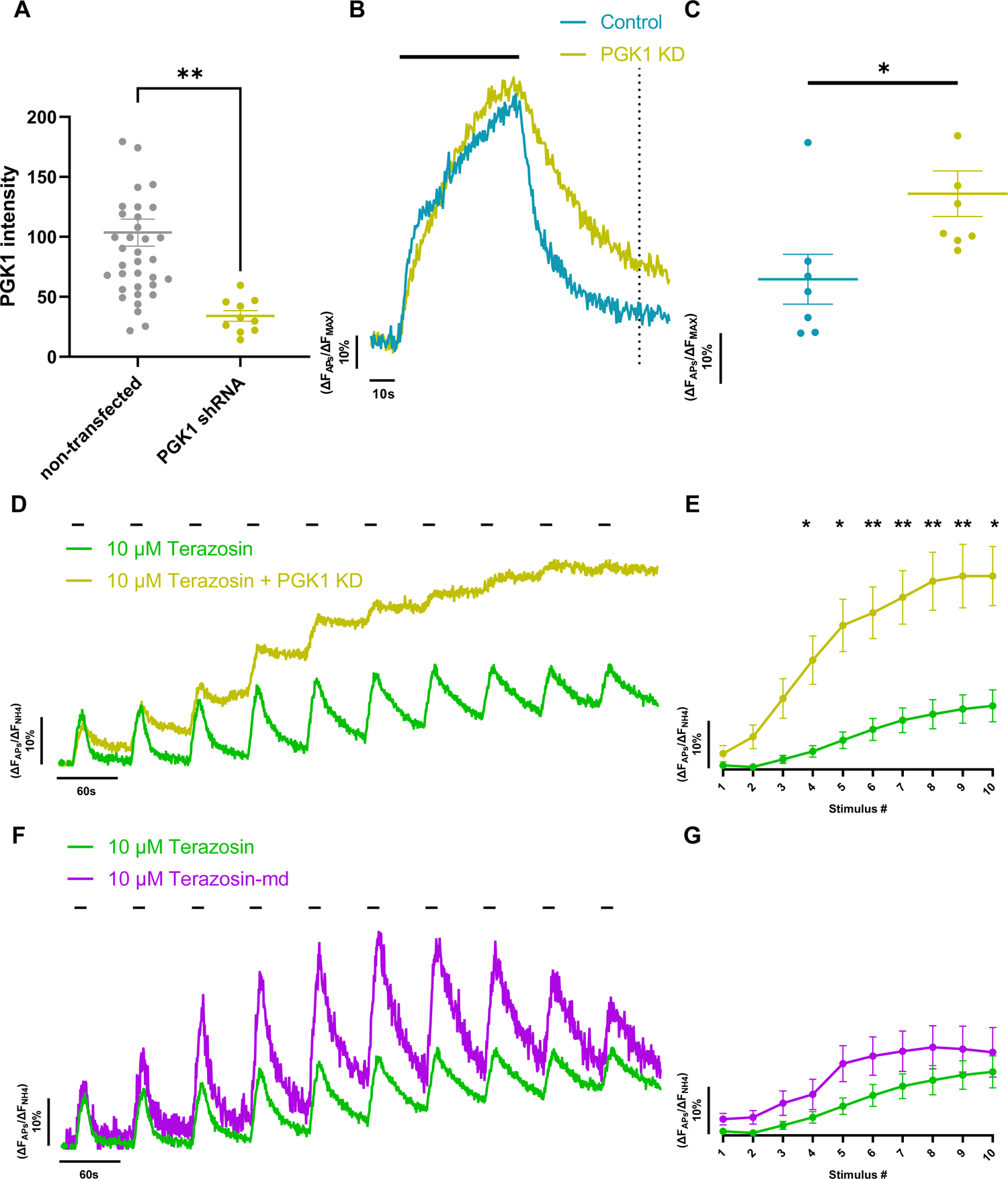
The terazosin effect is specific to PGK1. **(A)** Quantification of the KD efficiency of PGK1 from transfected cells with a PGK1 shRNA (yellow) and control (grey) and stained against PGK1. Mean ± SEM, non-transfected N=37, PGK1 shRNA N=10, \*\**p* < 0.01 unpaired t-test. (**B**) Ensemble average vGlutI-pH traces in 5 mM Glucose (teal) compared to PGK1 KD (yellow) stimulated with 600 APs (10 Hz, indicated by the black bar). (**C**) Quantification of the remaining fluorescence of the vGlut-pH traces 60 s post stimulation (indicated by the dotted line) mean ± SEM, wt N=7, wt PGK1 KD N=8, *p < 0.05 unpaired t-test. (**D**) KD of PGK1 (yellow) completely abolishes the Terazosin hypometabolic rescue (green). (**E**) Quantification of the remaining fluorescence from (**D**). mean ± SEM, 10 μM Terazosin N=10, 10 μM Terazosin + PGK1 KD N=14 *p < 0.05, **p < 0.01 2-way ANOVA. (**F**) Terazosin-md that binds PGK1 but not A1 (purple) still offers protection. (**G**) Quantification of the remaining fluorescence from (**F**). mean ± SEM, Control N=23, 10 μM Terazosin-md N=6.

**Fig. S3.**
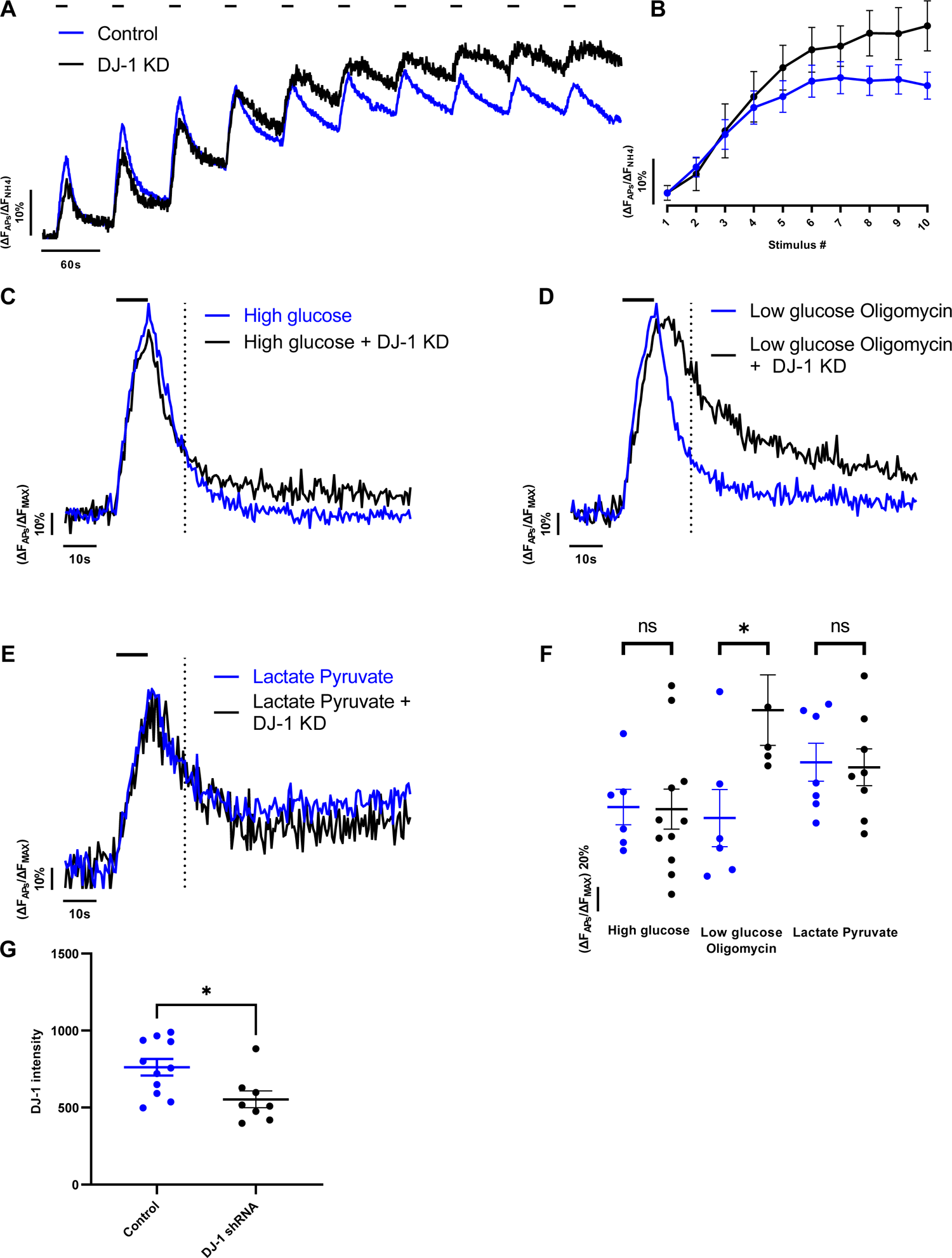
DJ-1 metabolic sensitivity. **(A)** DJ-1 KD (black) has only a mild phenotype in the synaptic endurance assay, also quantified in (**B**). mean ± SEM, Control N=23, DJ-1 KD N=6. (**C**) Ensemble average vGlutI-pH traces in response to a single 100 AP burst (10 Hz, indicate by black bar) are similar in control (blue) and DJ-1 KD neurons (black) in 5 mM glucose, but (**D**) are significantly slowed (blue and black respectively) with a combination of 0.1 mM glucose and 2 μM Oligomycin. (**E**) Ensemble average vGlutI-pH traces in response to a single 100 AP burst (10 Hz, indicate by black bar) are similar in control (blue) and DJ-1 KD neurons (black) in 1.25 mM Lactate and Pyruvate. (**F**) Fluorescence recovery 10 s post stimulus for the traces in (**C**, **D**, **E**) mean ± SEM, High glucose Control N=6, High glucose DJ-1 KD N=11, Low glucose Oligomycin Control N=6, Low glucose Oligomycin DJ-1 KD N=5, Lactate Pyruvate Control N=7, Lactate Pyruvate DJ-1 KD N=8,. p^ns^ > 0.05, *p < 0.05 unpaired t-test. (**G**) Quantification of the DJ-1 KD efficiency mean ± SEM, Control N=11, DJ-1 shRNA N=8, \**p* < 0.05 unpaired t-test.

**Fig. S4.**
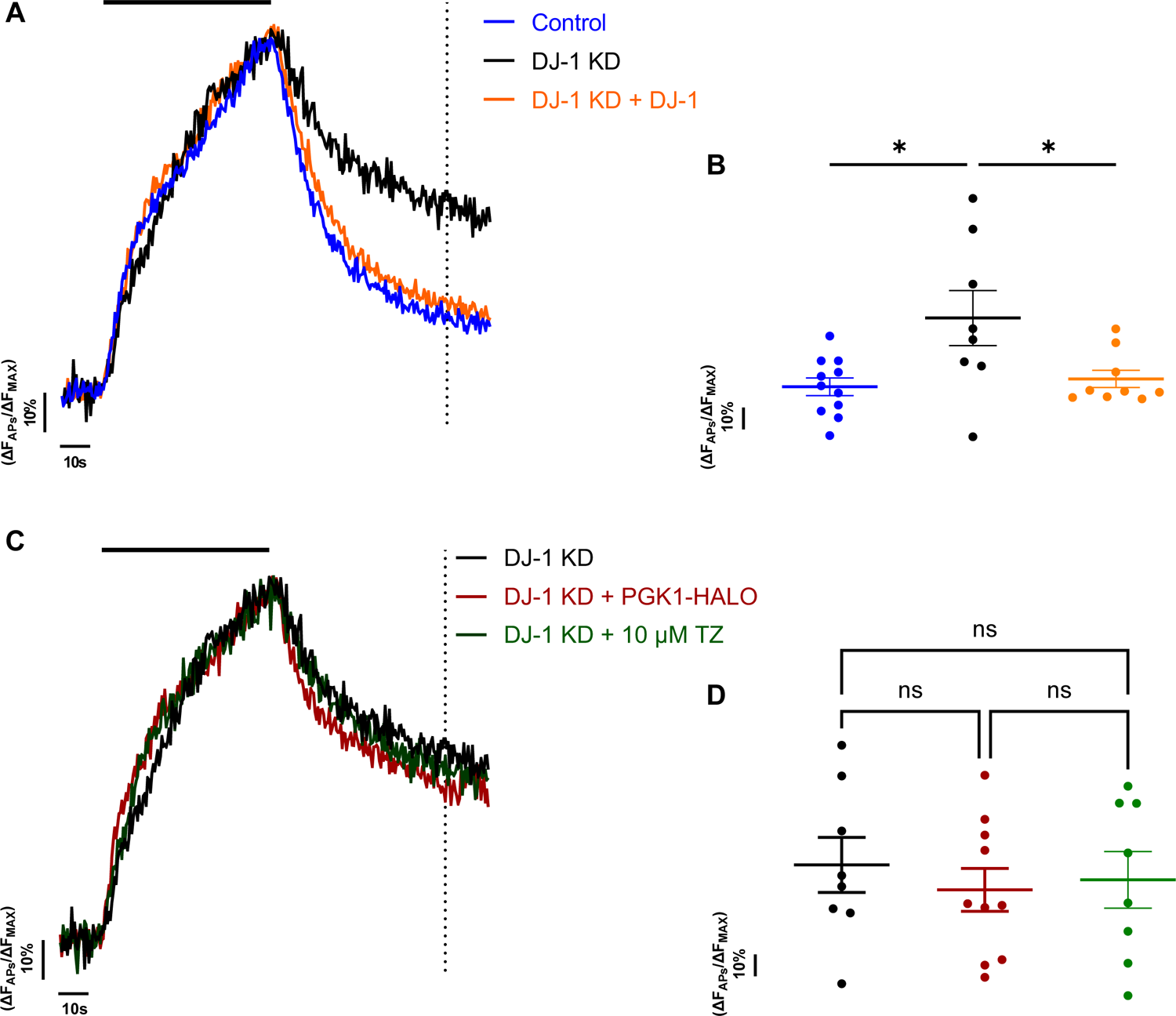
PGK1 activation cannot rescue the DJ-1 KD phenotypes. **(A)** DJ-1 KD (black) slows SV endocytosis in 0.1 mM glucose when cells were challenged with 600 APs at 10 Hz (black bar), compared to control (blue). Reintroduction of DJ-1 (orange) completely restores SV kinetics. (**B**) Quantification of the remaining fluorescence 60 s post stimulation (time point highlighted by dotted line in (**A**) mean ± SEM, Control N=11, DJ-1 KD N=8, DJ-1 KD + DJ-1 N=9 \**p* < 0.05 one-way ANOVA. (C) In the same conditions, upregulation of PGK1 (PGK1-HALO red, TZ green) is unable to restore SV kinetics. (**D**) Quantification of the remaining fluorescence mean ± SEM, DJ-1 KD N=8, DJ-1 KD + PGK1-HALO N=10, DJ-1 KD + 10 μM Terazosin N=8 n^s*p*^ > 0.05 one-way ANOVA.

